# Transcription factor-mediated generation of dopaminergic neurons from human iPSCs – a comparison of methods

**DOI:** 10.1101/2024.02.22.581485

**Authors:** Kirstin O. McDonald, Nikita M.A. Lyons, Lucia Schoderboeck, Stephanie M. Hughes, Indranil Basak

## Abstract

Dopaminergic neurons are the predominant brain cells affected in Parkinson’s disease. With the limited availability of live human brain dopaminergic neurons to study pathological mechanisms of Parkinson’s disease, dopaminergic neurons have been generated from human skin cell-derived induced pluripotent stem cells. Originally, induced pluripotent stem cell-derived dopaminergic neurons were generated using small molecules. These neurons took more than two months to mature. However, transcription factor-mediated differentiation of induced pluripotent stem cells has revealed quicker and cheaper methods to generate dopaminergic neurons. In this study, we compare and contrast three protocols to generate induced pluripotent stem cell-derived dopaminergic neurons using transcription factor-mediated directed differentiation. We deviated from the established protocols using lentivirus transduction to stably integrate transcription factors into induced pluripotent stem cells, followed by differentiation using different media compositions. We introduced three transcription factors into the AAVS1 safe harbour locus of induced pluripotent stem cells, and in combination with small molecules, we generated more than 80% neurons in the culture, out of which more than 80% neurons were dopaminergic neurons. Therefore, a combination of transcription factors along with small molecule treatment may be required to generate a pure population of human dopaminergic neurons, a prerequisite for cell replacement therapies.

## 1. Introduction

Induced pluripotent stem cell (iPSC) research has helped neuroscientists minimise the dependence on transgenic/knockout animal models and human cells like embryonic stem cells or cells derived from biopsies [1, 2]. The versatility of iPSCs has been harnessed in other fields of research, such as cardiac and renal diseases [3, 4]. Many groups have embarked on differentiating skin fibroblast-derived iPSCs into dopaminergic neurons, from both healthy and Parkinson’s disease affected individuals, using different approaches such as small molecules, transcriptions factors, and microRNAs [1, 5–13]. The original protocol published more than a decade ago by Kriks *et al*. [6] used small molecules to generate human dopaminergic neurons from iPSCs, however, the protocol took more than two months to achieve mature human dopaminergic neurons. Other studies [14–16] have used modified protocols with small molecules and have obtained purer populations of dopaminergic neurons, however, the time to develop the mature neurons was still more than a month. Since 2013, several researchers have developed protocols for transcription factor-mediated differentiation, using various transcription factors [7–11], and have successfully generated iPSC-derived human dopaminergic neurons in a relatively short amount of time (less than one month). Theka *et al*. [5] was one of the first to generate iPSC-derived dopaminergic neurons through transcription factor-mediated differentiation, omitting the neural progenitor cell stage. The protocol involved lentivirus transduction of four components - reverse tetracycline-controlled transactivator (rtTA) and three transcription factors, Ascl1, Nurr1 and Lmx1a. Nishimura *et al*. 2022 [8] used a PiggyBac Transposase Expression Vector system to integrate two transcription factors (ASLC1 and LMX1A) to generate human dopaminergic neurons. In the same year, Sheta *et al* [9] used only one transcription factor (NGN2) to differentiate iPSCs into dopaminergic neurons. However, the bottleneck has always been getting a pure population of dopaminergic neurons, something that Zhang *et al* [17] and Fernandopulle *et al*. [18] achieved with human glutamatergic cortical-like neurons. This bottleneck is a hindrance to cell replacement therapy in Parkinson’s disease, where dopaminergic neurons are the most vulnerable neuronal population in the brain and the first ones to degenerate.

In this study, we modified and compared three different protocols using transcription factors to generate human dopaminergic neurons. We stably integrated transcription factors into iPSCs using CRISPR-Cas9 editing and controlled their expression through a tetracycline-inducible promoter. The stable integration enabled us to genetically modify the iPSCs in a one-step protocol and eliminated variation from co-transduction or sequential transduction of different lentiviruses. CRISPR-Cas9 gene editing causes double-strand breaks, which also induces a p53-mediated cell death [19]. To counter this problem, we used a non-integrating dominant negative p53 construct along with the donor DNA (transcription factors), single guide RNAs and Cas9 protein while generating the iPSC lines stably expressing the transcription factors.

Using different media compositions, we achieved the generation of iPSC-derived human dopaminergic neurons (iDA) from all three protocols within three weeks. However, the percentage of iDAs varied significantly from what has been reported by the respective studies. The modification we applied to the protocol from Sheta *et al*. yielded the highest percentage of tyrosine hydroxylase and GIRK2 positive dopaminergic neurons in 3 weeks, which were also electro-physiologically active. Our results suggest a finely optimised combined approach may be required for the generation of pure populations of dopaminergic neurons for transplantation purposes.

## 2. Materials and Methods

### 2.1 Reagents

For the three different protocols to generate iPSC-derived iDAs, the following reagents were used. Media compositions are included in Table S3.

For iPSCs – WTC11 human induced pluripotent stem cells (gifted by Dr Michael Ward, National Institute of Health (NIH), Bethesda, MD, USA), Matrigel (cat# 354277), Rock Inhibitor (RI) Y-27632 (cat# RDS125410) (both from In Vitro Technologies, Auckland, NZ), Essential 8 (E8) medium (cat# A1517001), EDTA (0.5M) pH 8.0 (cat# AM9261), PBS pH

7.4 (cat# 70011044), Accutase (cat# A1110501) (all from ThermoFisher Scientific, Auckland, NZ).

For genotyping and polymerase chain reactions - PureLink Genomic DNA kit (cat# K182001, ThermoFisher Scientific, Auckland, NZ), Phusion High-Fidelity DNA Polymerase (cat# M0530L, New England Biolabs, MA, USA), and Sanger sequencing at Massey Genome Service, Palmerston North, NZ.

For recombinant lentivirus production - HEK293FT cell line (cat# R70007, ThermoFisher Scientific, Auckland, NZ), HT1080 cell line (cat# CCL-121, ATCC Manassas, VA, USA), psPAX2 (cat# 12260, Addgene, Watertown, MA, USA), VSVg (cat# K497500), DMEM, high glucose, pyruvate, no glutamine (cat# 10313021), MEM Non-Essential Amino Acids Solution (100X) (cat# 11140050), L-Glutamine (200 mM) (cat# 25030081), Sodium Pyruvate (100 mM) (cat# 11360070), Penicillin-Streptomycin (10,000 U/mL) (cat# 15140122), Opti-MEM™ Reduced Serum Medium (cat# 31985070), TrypLE™ Express Enzyme (1X), phenol red (cat# 12605028), Lipofectamine™ 2000 Transfection Reagent (cat# 11668019) (all from ThermoFisher Scientific, Auckland, NZ), Fetal Bovine Serum – Sterile Filtered (cat# FBSF, (Moregate BioTech, Bulimba, QLD, Australia), Polybrene (H9-268), Poly-L-Lysine hydrochloride (cat# 9404), Lactose (cat# MR5245) (all from Merck, Auckland, NZ).

For iPSC-derived iDA generation following Theka *et al*. [5] – Alt-R CRISPR-Cas9 AAVS1 sgRNAs and Alt-R S.p. Hifi Cas9 Nuclease V3 (cat# 104906119) (IDT, Melbourne, VIC, Australia), OptiMEM I redcued serum medium (cat# 31985070), Lipofectamine Stem transfection reagent (cat# STEM00001), DMEM/F12 (cat# 11330032), Penicillin-Streptomycin (10,000 U/mL) (cat# 15140122), B27 supplement (cat# 17504044) (all from ThermoFisher Scientific, Auckland, NZ), puromycin dihydrochloride (cat# P8833), insulin (cat# I9278), transferrin (cat# T8158), sodium selenite (cat# 214485), progesterone (cat# P6149), putrescine (cat# P5780) doxycycline (cat# D9891), paraformaldehyde (cat# P6148), and 4′,6-diamidino-2-phenylindole (DAPI) (D9542) (all from Merck, Auckland, NZ). See table S2 for antibodies used for immunocytochemistry (ICC).

For iPSC-derived iDA generation following Nishimura *et al*. [8] - Essential 6 medium (E6) (cat# A1516401), Neurobasal medium (cat# 21103049), B27 supplement minus vitamin A (cat# 12587010), GlutaMax (cat# 35050061) (all from ThermoFisher Scientific, Auckland, NZ), puromycin dihydrochloride (cat# P8833), doxycycline (cat# D9891), dibutyryl cyclic AMP (dbcAMP; cat# D0627) (all from Merck, Auckland, NZ), DAPT (cat# 2088055, Peprotech/Lonza, Sydney, Australia), brain-derived neurotrophic factor (BDNF; cat# RDS248BDB050, In Vitro Technologies, Auckland, NZ), glial cell derived neurotrophic factor (GDNF; cat# 450-01), and ascorbic acid (cat# 5088177) (both from PeproTech, Rocky Hill, NJ, USA). See table S2 for antibodies used for ICC.

For iPSC-derived iDA generation following Sheta *et al*. [9] - DMEM/F12 (cat# 11330032), L-Glutamine (cat# 25030081), Non-essential amino acids (NEAA; cat# 11140050), N-2 supplement (cat# 17502048), laminin (cat# 23017015), B27 supplement (cat# 17504044) (all from ThermoFisher Scientific, Auckland, NZ), Poly-L-Ornithine (cat# P3655), Cytosine β-D-arabinofuranoside hydrochloride (Ara-C) (cat# C6645) (both from Merck, Auckland, NZ). BDNF (cat# RDS248BDB050), NT-3 (cat# RDS267N3025) (both from In Vitro Technologies, Auckland, NZ), BrainPhys Neuronal Media (cat# 05790), STEMdiff Midbrain Neuron Differentiation kit (cat# 100-0038), STEMdiff Midbrain Neuron Maturation kit (cat# 100-0041), human recombinant shh (C24II) (cat# 78065.1) (all from STEMCELL Technologies, Tullamarine, VIC, Australia). See table S2 for antibodies used for ICC.

### 2.2 iPSC-derived dopaminergic neuron generation following Theka et al

Theka *et al*. [5] presented the first data showing that the expression of three transcription factors, ASCL1, NURR1 and LMX1A, can drive iPSCs towards dopaminergic neurons (iDAs). The authors used lentiviral methods to induce the expression of the three transcription factors into iPSCs followed by use of a neuronal inducing media for 7-18 days to generate dopaminergic neurons. We used the two following different methods to integrate the transcription factors into the iPSCs.

#### 2.2.1 Overexpression of transcription factors using recombinant lentiviruses

The transcription factors and the reverse tetracycline transactivator (rtTA) were obtained from Addgene (Tet-O-Fuw-Ascl1 Addgene plasmid #27150, pLV.PGK.mLmx1a Addgene plasmid #33013, Nurr1 Addgene plasmid #35000, FUW-M2rtTA Addgene plasmid #20342). The cassette with the three human transcription factors hALAN – <u>h</u>uman <u>A</u>SCL1, <u>L</u>MX1<u>A</u> and <u>N</u>URR1 was synthesized by GeneArt Gene Synthesis Service (ThermoFisher Scientific, Auckland, NZ) and cloned into a lentiviral plasmid with mNeonGreen as a marker driven by EF1a promoter (Fig. S1A). The rtTA was cloned into another lentiviral plasmid with blue fluorescence protein (BFP) as a marker driven by EF1a promoter (Fig. S1A). Lentiviral generation was performed following our established protocol [20]. iPSCs were transduced with the transcription factor lentivirus and the rtTA lentivirus at a multiplicity of infection (MOI) of 1, followed by fluorescence-activated cell sorting (FACS) [21] for mNeonGreen and BFP (Fig. S1B). The double positive (mNeonGreen and BFP) enriched iPSCs were expanded and cryopreserved for downstream differentiation into iDAs.

#### 2.2.2 Integration of transcription factors using CRISPR-Cas9 approach

The synthesized hALAN was cloned into a donor construct to introduce the hALAN into the AAVS1 safe harbour site for iPSC differentiation (Addgene plasmid #105840). The resulting plasmid expresses hALAN, rtTA and mApple (Fig. 1A). To stably integrate the hALAN cassette into the AAVS1 site and generate iPSCs expressing hALAN for iDA differentiation, a workflow outlined in Fig. S2A-D was followed. iPSCs were transfected with 500 ng of the plasmid, 3 pmol of 2 sgRNAs against AAVS1 (Table S1 for sgRNA sequences), 3 pmol of Alt-R HiFi Cas9 enzyme and 100 ng of pCE-mp53DD (dominant-negative p53; addgene plasmid # 41856) using Lipofectamine Stem following the manufacturer’s protocol. The day following transfection, transfected iPSCs were passaged with Accutase and plated for expansion. The mApple positive iPSCs were isolated by FACS, and the sorted iPSCs were expanded, cryopreserved and plated for clonal outgrowth at 50% serial dilutions. Individual colonies were picked manually, expanded and treated with 1 μg/mL puromycin followed by genotyping to ensure the integration of hALAN.

**Figure 1.**
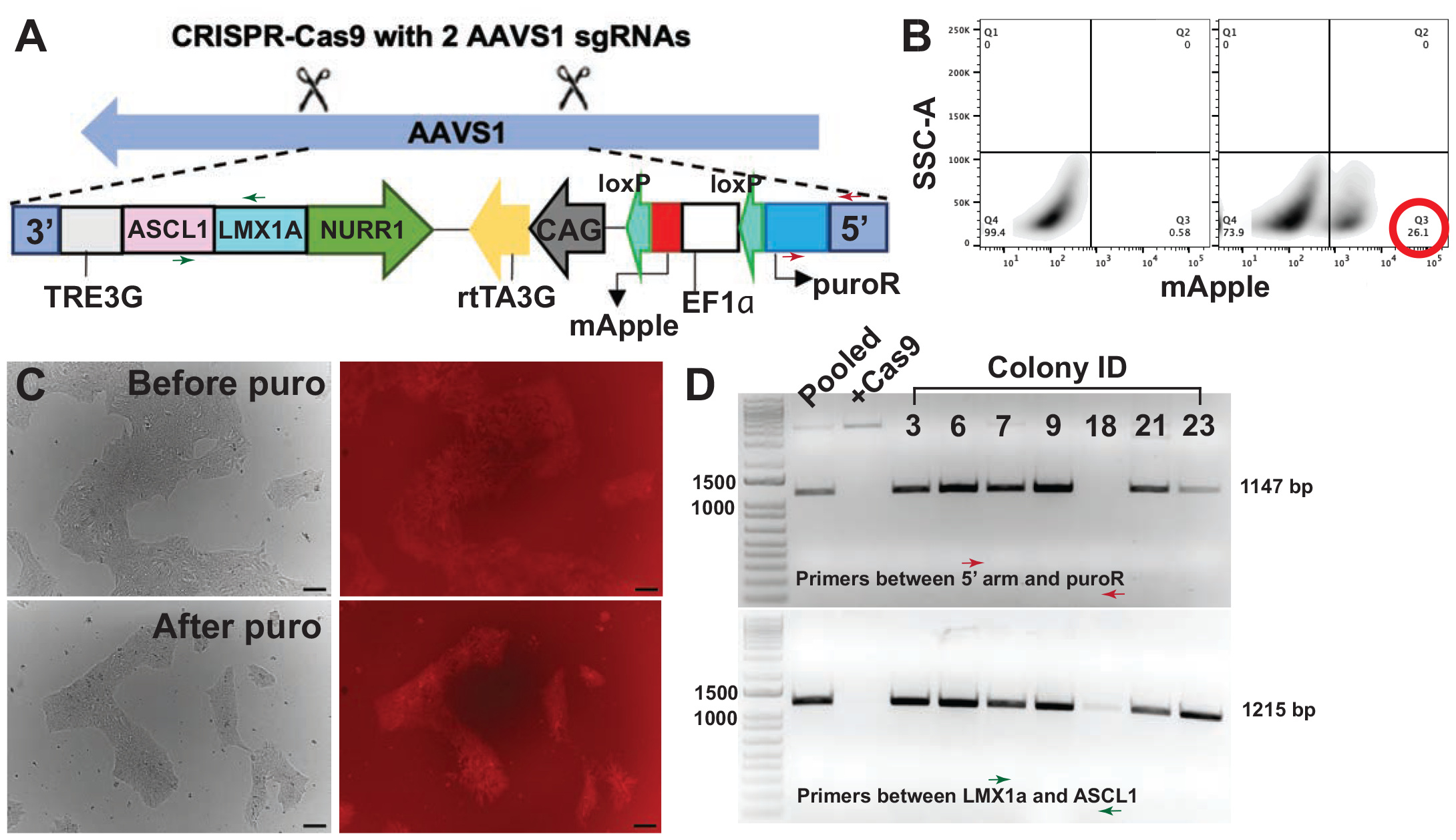
Integration of transcription factors and selection of iPSC colonies. **A**. Using CRISPR-Cas9, the cassette containing the 3 human transcription factors (ASCL1, LMX1A and NURR1), puroR gene, and the mApple selection marker were integrated into the AAVS1 safe harbour locus of iPSCs. **B**. FACS was used to isolate the mApple positive iPSCs (> 25 %) post CRISPR-Cas9 edit of the iPSCs with the hALAN cassette. **C**. Post-FACS, iPSC colonies were manually picked and treated with puromycin to obtain a purer population of mApple positive iPSCs. Scale bar 100 μm. **D**. The FACS and puromycin-treated iPSC colonies were genotyped using PCR amplifying a partial region of the 5’ arm and the puroR (red arrows) and a partial region of the LMX1a and the ASCL1 transcription factors (green arrows). Also refer to **A** for the position of the primers indicated by the red and the green arrows. DNA gel showing ladder in lane 1, pooled iPSCs (before colony picking) in lane 2, Cas9 only iPSCs (no hALAN integration) in lane 3 and colonies with positive/negative PCR bands in the following lanes. The numbers above the lanes represent colony ID numbers.

#### 2.2.3 Genotyping of iPSCs with integrated hALAN

Genomic DNA was extracted from the mApple-sorted iPSC colonies using PureLink Genomic DNA kit following the manufacturer’s protocol. Genomic DNA was quantified using Nanodrop and used for PCR with Phusion polymerase following the manufacturer’s protocol and Table S1) to detect the transcription factors. Primers used for genotyping are included in Table S1.

#### 2.2.4 Differentiation of iPSCs into iDAs

Following the original protocol published by Theka *et al*. [5], the mApple sorted iPSCs with the three transcription factors integrated at the AAVS1 site were differentiated using the media composition outlined in Fig. S2E. iPSCs were passaged with Accutase and plated on Matrigel coated plate in E8+RI. After two days, the E8 media was replaced with neuronal inducing medium containing DMEM/F12, 25 µg ml^-1^ insulin, 50 µg ml^-1^ transferrin, 30 nM sodium selenite, 20 nM progesterone, 100 nM putrescine and penicillin/streptomycin with the addition of doxycycline (2 μg/ml). After 7 days of differentiation, B27 was added to the aforementioned media and kept for the following 7-18 days. The medium was changed every 2-3 days and the cells were maintained in culture medium for 21 and 28 days for the maturation of the iDAs followed by assessment by ICC.

### 2.3 iPSC-derived dopaminergic neuron generation following Nishimura et al

The following protocol was adopted from Nishimura *et al*. [8] to differentiate the hALAN integrated mApple sorted iPSCs into iDAs (Fig S3A). iPSCs were passaged with Accutase and plated on Matrigel coated plate in E8+RI. Two days after plating, the E8 media was replaced with Essential 6 medium (E6) supplemented with 5.0 mg/mL DOX and 1.0 mg/mL puromycin for the first 3 days. After 3 days, the E6 medium was changed to Neurobasal medium supplemented with B27 supplement vitamin A minus, GlutaMax, 5.0 mg/mL DOX, and 1.0 mg/mL puromycin and maintained for the next 4 days. After three days in E6 media and 4 days in Neurobasal media, the neuronal inducing media was supplemented with 10 mM DAPT for the next 3 days followed by supplementing with 20 ng/mL BDNF, 10 ng/mL GDNF, 200 mM ascorbic acid, and 500 mM dbcAMP and continued until day 21. On day 21, the iDAs were used for ICC.

### 2.4 iPSC-derived dopaminergic neuron generation following Sheta et al

The following protocol was adopted from Sheta *et al*. [9] to differentiate the iPSCs into iDAs. With an established protocol to generate iPSC-derived cortical neurons (i^3^Ns) [18, 22], our i^3^N protocol was combined with the iDA protocol from Sheta *et al*. to modify the media compositions and generate purer populations of iDAs. For this protocol, the hALAN iPSCs were not used, rather iPSCs with integrated NGN2 transcription factor were used [17, 18, 22]. These NGN2-iPSCs express NGN2 under the control of a doxycycline-inducible promoter, as developed by Fernandopulle *et al*. [18]. The NGN2-integrated iPSCs were initially plated on a Matrigel-coated plate in E8+RI, and after two days of culture (ensuring colony formation), the iPSCs were passaged with Accutase to get a single-cell suspension. The single cell suspension of iPSCs was re-plated on a Matrigel-coated plate in warm Induction media (Table S3) supplemented with doxycycline (2 µg/mL) and RI (1 µg/mL). For the next two days, the Induction media was replaced with more warm Induction media supplemented with doxycycline (2 µg/mL). After three days of doxycycline-mediated induction of NGN2 expression, the induced iPSCs were harvested using Accutase and re-plated on poly-L-ornithine (PLO, 100 μg/mL final concentration) and laminin (10 µg/mL final concentration)-coated plates in cortical neuron media (Table S3). Two days after plating the induced iPSCs in cortical neuron media, the cortical neuron media was completely replaced with STEMDiff Midbrain Differentiation Media (Table S3) supplemented with Cytosine β-D-arabinofuranoside hydrochloride (Ara-C) at 2 µM (to inhibit the proliferation of any remaining dividing cells) and doxycycline (2 μg/mL). The differentiation media was completely replaced with fresh iDA Differentiation Media supplemented with Ara-C at 2 µM and doxycycline (2 μg/mL) for the next two days followed by half media changes (iDA differentiation Media supplemented doxycycline (2 μg/mL)) for another four days. Finally, after eight days of differentiation, the media was completely replaced with fresh iDA Maturation Media (Table S3) supplemented with doxycycline (2 μg/mL) on the 9^th^ day. Starting from day 10 and every two days, half of the media was replaced with fresh iDA Maturation Media supplemented with doxycycline (2 μg/mL) for 22 and 26 days for the maturation of the iDAs followed by assessment by ICC.

### 2.5 Immunocytochemistry and calcium signalling on iPSC-derived iDAs

The iPSC-derived iDAs from all three protocols were used for ICC to ascertain the percentage of dopaminergic neurons being generated. ICC was performed following our previously established protocol [22]. Briefly, the iPSC-derived iDAs were maintained for 3-4 weeks, in different media conditions as outlined in the previous three sections and were fixed using 4% paraformaldehyde in PBS. The neurons were stained with antibodies against TUJ1 (neuron-specific class III beta-tubulin), MAP2 (dendritic neuronal marker), tyrosine hydroxylase (dopaminergic neuronal marker), GIRK2 (G-protein-regulated inward-rectifier potassium channel 2, neuronal marker characteristic of dopaminergic neurons from substantia nigra of human brain), LMX1A and NURR1 (transcription factors used to generate iPSC-derived dopaminergic neurons (Table S3). Alexa Fluor-labelled secondary antibodies (ThermoFisher) were used as per Table S3 followed by staining with 4′,6-diamidino-2-phenylindole (DAPI). The stained neurons were imaged on a Nikon Eclipse Ti2 epifluorescence microscope (Nikon, Tochigi, Japan) and Olympus FV3000 confocal microscope (Olympus, Tokyo, Japan). For calcium imaging, i^3^Ns and iDAs were transduced with lentivirus expressing GCaMP7s driven by the Synapsin promoter (pCDH.rSyn.jGCaMP7s-cloned from pGP-AAV-syn-jGCaMP7s-WPRE, Addgene plasmid #104487) two days before live cell imaging. Using an Olympus FV3000 confocal microscope (Olympus, Tokyo, Japan) with a 37 °C heated chamber, 100 frames were captured per field (n ≥ 3 fields per condition) with an interval of 2 second to obtain a time lapse video of calcium signaling in the i^3^Ns and the iDAs.

### 2.6 Image analysis and statistical analysis

Images acquired on the Nikon and Olympus microscopes were analysed using ImageJ (NIH, Bethesda, MD, USA). For DAPI counts, the following workflow was used. Open (image) > Image > Adjust Threshold (set threshold to separate the individual DAPI positive nuclei, in this case it was set between 0-6600) > Process > Binary > Make binary > Process > Binary > Outline > Measure > Analyse Particle > Set Size (0.50-infinity) and Circularity (0.00-1.00) > Show Outlines > Note the ‘Count’ from the Summary window. For TUJ1, MAP2 and TH counts, the neuronal cell bodies were counted showing positive staining using a similar workflow as above. The counts for TUJ1, MAP2 and TH positive cells were normalized by the counts for DAPI followed by calculating percentage of TUJ1, MAP2 and TH positive cells on MS Excel. The percentage positive values were imported onto GraphPad Prism (GraphPad, San Diego, CA, USA), tested for normal distribution and unpaired two-tailed student t-tests were performed to assess statistical significance, reported as p values. Data are presented as mean ± standard error of mean (SEM). * indicates p < 0.05.

## 3. Results

### 3.1 Targeted and stable integration in iPSCs maintained transcription factor expression better than lentiviral transduction

Post-transduction of the two lentiviruses (Fig. S1A) in the iPSCs, following Theka *et al* [5], the iPSCs were sorted for BFP and mNeonGreen (Fig. S1B). The percentage of double positive iPSCs (BFP+, mNeonGreen +) was less than 15% (Fig. S1B, second plot) and the double positive iPSCs were expanded (Fig. S1Ci) for differentiation. However, with passaging, the double positive iPSCs started losing the mNeonGreen fluorescence (Fig. S1Cii). Although whether the hALAN construct was lost from the iPSCs was not verified, with the recent success with stably integrating transcription factors into iPSC safe harbour loci [18], an alternative method (Fig. S2) to integrate hALAN was adopted. The alternative method involved transfection of the hALAN cassette (Fig. S2A), FACS (Fig. S2B), serial dilution (Fig. S2C), manual picking and puromycin selection, and validation of clones by genotyping (Fig. S2D). Using CRISPR-Cas9, the hALAN construct along with the mApple, puroR and rtTA, were integrated into the AAVS1 safe harbour locus of iPSCs (Fig. 1A), which yielded mApple+ iPSCs (Fig. 1B, second plot). The bulk FACS of mApple yielded > 25% mApple+ iPSCs (Fig. 1B, second plot), which were expanded, cryopreserved and later re-plated to pick colonies manually. The picked colonies were treated with puromycin to obtain a purer population of mApple+ iPSCs (Fig. 1C) followed by genotyping by PCR (Fig. 1D) and sequencing. Using primers spanning ASCL1 + LMX1A and 5’ AAVS1 arm + PuroR, 6 colonies were identified that had hALAN integrated into the AAVS1 site. Colonies #6 and #9 were used for the differentiation protocol.

### 3.2 Differentiation following Theka et al and Nishimura et al

Fig. S2E summarises the differentiation protocol with different media compositions over 3 weeks adapted from Theka et al. Similar to the original paper, our ICC experiments after 3 weeks of differentiation showed the generation of TUJ1+ neurons. However, day 21 cells showed only 5% TUJ1+ neurons (Fig. 2A, C) and only 2% TH staining (Fig. 2A, white arrow showing weak TH staining in neuronal processes). When the differentiation was extended to 28 days, the percentage of TUJ1+ cells improved significantly to 13% (Fig. 2B, C), but still showed low TH staining (4%) (Fig. 2B, white arrow showing weak TH staining in neuronal processes). TH staining was brighter in the cell bodies compared to the neuronal processes (Fig. 2A, B). Combining the percentage of TH+ and TUJ1+ cells, day 21 cells showed 37% of the TUJ1+ cells were also TH+, whereas day 28 cells showed 28% of the TUJ1+ cells were also TH+ (Fig. 2C). There was no significant difference between day 21 and day 28 percentage of TH+/TUJ1+ neurons (Fig. 2C). Our i^3^N protocol [22] uses PLO instead of Matrigel to coat cell culture plates for plating and differentiating neurons. Therefore, as an alternative coating surface, PLO-coated wells were also used in addition to Matrigel-coated wells. As iPSCs would not survive on PLO-coated wells, the differentiation was started following Theka *et al* [5], and after 7 days of differentiation (before switching to B27 media on Day 8), pre-differentiated cells were lifted with Accutase and plated on PLO-coated wells and maintained until day 21 or 28. Our modified protocol on PLO-coated wells did not improve the generation of TUJ1+ neurons (Fig. S2F, G). Therefore, we further modified the existing protocol by incorporating new media compositions (Fig. S3A) following Nishimura *et al*[8]. ICC on the iDAs differentiated from hALAN iPSCs using the Nishimura protocol resulted in improved generation of TUJ1+ neurons (74%) (Fig. 2D, E). More than 22% of the cells also stained positive for TH (Fig. 2D, E). However, only 31.2% of TUJ1+ neurons were also TH+, indicating that the population of neurons being generated from this protocol are predominantly not TH+, i.e., not dopaminergic neurons (Fig. 2D, E, Fig. S3B). GIRK2 is a marker characteristic of dopaminergic neurons from the substantia nigra and the central tegmental area [23] and hence, we also tested for GIRK2 expression in the iPSC-derived iDAs alongside expression of the transcription factors LMX1A and NURR1 that are involved in dopaminergic neurogenesis (also present in the hALAN cassette). Interestingly, most cells showed GIRK2+ staining (compare overlap of GIRK2 and DAPI in Fig. S3C), but not all cells showed a distinct expression of LMX1A and NURR1 (compare overlap of LMX1A, NURR1 and DAPI in Fig. S3D). Therefore, although the Nishimura protocol showed a better yield of neurons, particularly dopaminergic neurons, the protocol produced an impure population of dopaminergic neurons with the possible inclusion of other types of unclassified cells.

**Figure 2.**
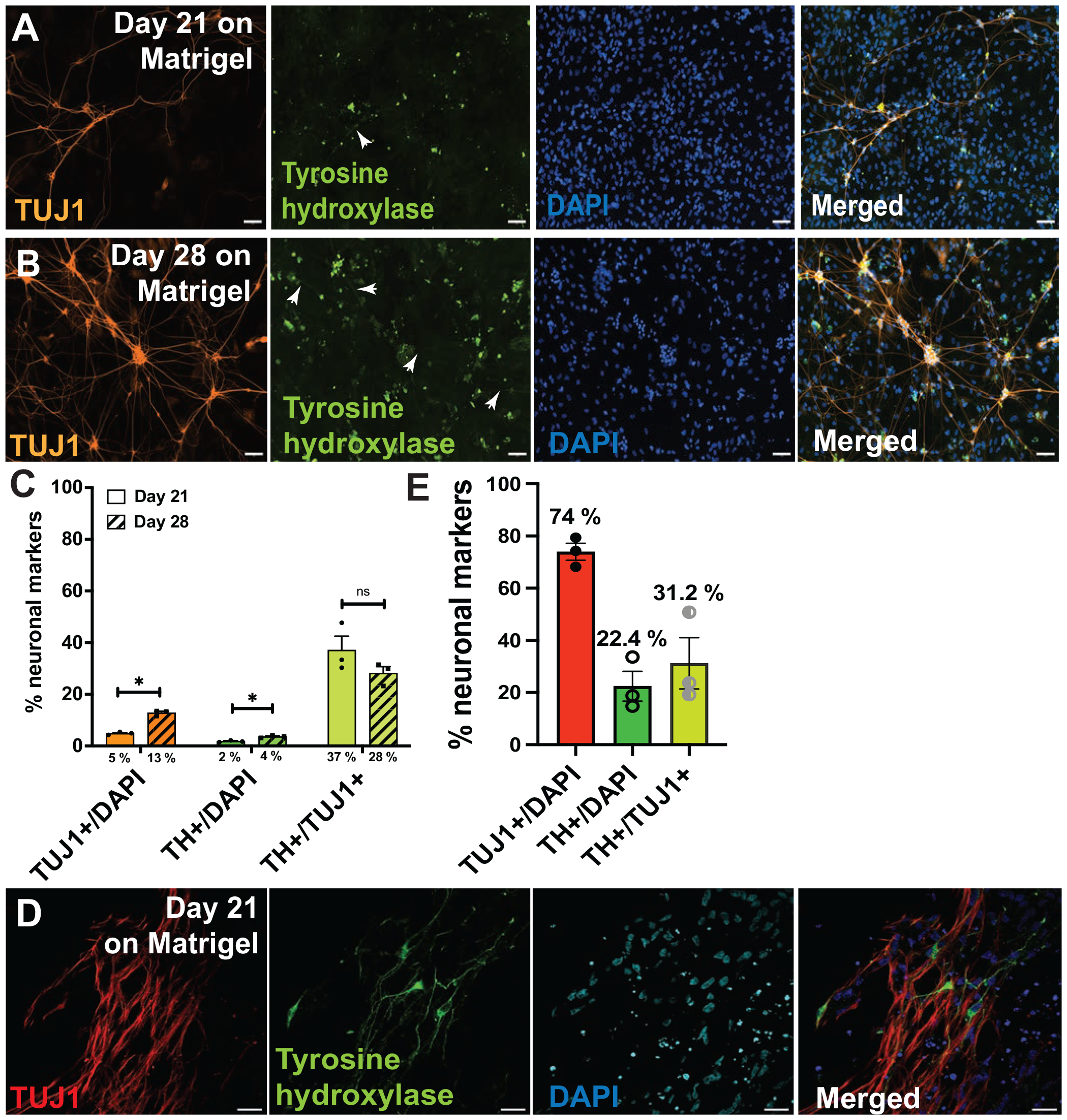
Differentiation of iPSCs into iDAs following Theka *et al*. and Nishimura *et al*. **A** and **B**. iPSC-derived day 21 and day 28 iDAs, respectively, following Theka *et al*. Cells were stained with TUJ1 (class III β tubulin, neuronal marker), tyrosine hydroxylase (dopaminergic neuronal marker) and DAPI (nuclear stain). Scale bar 100 μm. **C**. Quantification of TUJ1+ (over DAPI+), TH+ (over DAPI+) and TUJ1+ cells that are also TH+ in day 21 and day 28 iDAs following Theka et al. Percentages show the proportion of cells positive for respective markers. **D**. iPSC-derived day 21 iDAs, following Nishimura et al. Cells were stained with TUJ1 (class III β tubulin neuronal marker), tyrosine hydroxylase (dopaminergic neuronal marker) and DAPI (nuclear stain). Scale bar 20 μm. **E**. Quantification of TUJ1+ (over DAPI+), TH+ (over DAPI+) and TUJ1+ cells that are also TH+ in day 21 iDAs following Nishimura et al. Percentages show the proportion of cells positive for respective markers. Data are presented as mean ± standard error of mean. * *p* < 0.05, ns is *not significant*.

### 3.3 Differentiation following Sheta et al

Fig. 3A summarises the differentiation protocol with different media compositions over three weeks following Sheta *et al* [9]. The original protocol was modified to minimize the number of undifferentiated cells. By incorporating an Induction step (from Fernandopulle *et al* [18]), the iPSCs were pre-differentiated for 3 days before re-plating them at the desired number for downstream analysis (Fig. 3A). Twenty-one days post-differentiation, 77% of the cells were MAP2+, 63% of total cells were TH+, and 84% of the MAP2+ cells were TH+ (Fig. 3B, C). There was no staining for vGLUT1 (glutamatergic neuronal marker, Fig. 3B). This is the first time, to our knowledge, that over 80 % dopaminergic neurons have been obtained from iPSCs through transcription factor-mediated differentiation. When the protocol was extended to nearly 4 weeks, the MAP2+ staining increased to 84%. However, there was a significant drop in TH+ cells (44% on day 26 compared to 63% on day 21) and MAP2+ cells that were TH+ (53% on day 26 compared to 84% on day 21) (Fig. 3B, C). Co-staining for TH (63%) and GIRK2 (70%) showed even purer population of dopaminergic neurons at 95% (positive for both TH and GIRK2) (Fig. 3D, E). As a negative control, some pre-differentiated cells (gone through the induction step) were plated in cortical neuron media (Table S2) instead of STEMdiff media (Table S2), which stained positive for vGLUT1 and MAP2, but not for TH (Fig. S3E). Therefore, combining Fernandopulle and Sheta protocols yielded more than 84% of TH+/TUJ1+ dopaminergic neurons and 95% of the neurons were positive for both the dopaminergic markers TH and GIRK2. Furthermore, to test whether the iDAs are functional like the i^3^Ns[24], we expressed GCaMP7s calcium sensor via lentiviral transduction in three-week-old iDAs and i^3^Ns and recorded changes in calcium signalling as a surrogate for endogenous, non-stimulated electrophysiological properties. From our GCaMP7s experiment, it was observed that the i^3^Ns (Video S1A) iDAs are electrophysiologically active (Video S1A), similar to what was shown by Mahajani et al[11], and fire less spontaneously than the i^3^Ns (Video S1B).

**Figure 3.**
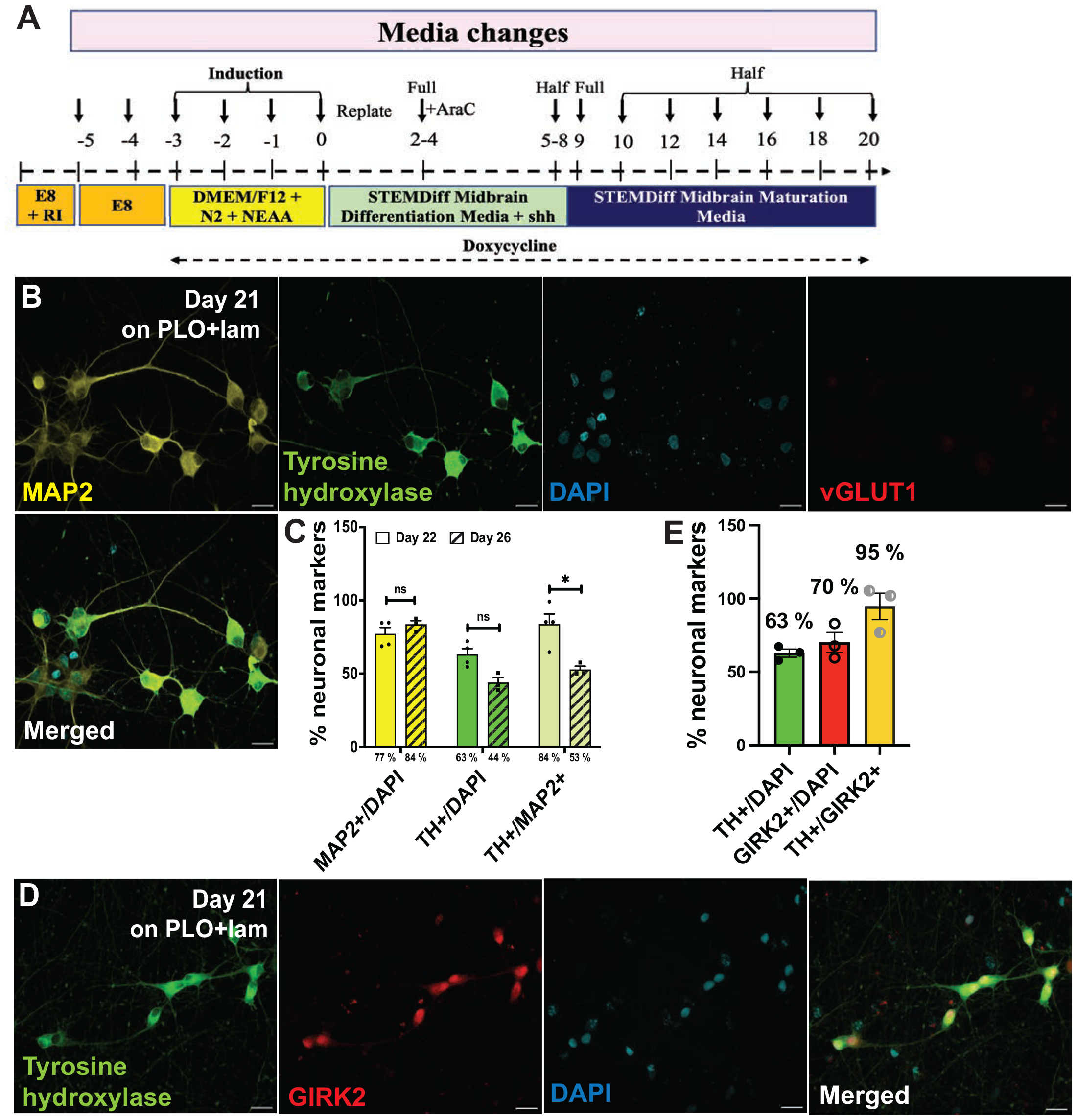
Differentiation of iPSCs into iDAs following Sheta *et al*. **A**. Schematic of the modified iPSCs-derived iDA protocol with media composition adapted from Sheta et al. An induction step was introduced into the original protocol published by Sheta et al. **B**. iPSC-derived day 22 iDAs following Sheta et al. Cells were stained with MAP2 (microtubule-associated protein, 2 neuronal marker), tyrosine hydroxylase (dopaminergic neuronal marker), vGLUT1 (vesicular glutamate transporter 1, glutamatergic neuronal marker) and DAPI (nuclear stain). Scale bar 20 μm. **C**. Quantification of TUJ1+ (over DAPI+), TH+ (over DAPI+) and TUJ1+ cells that are also TH+ cells in day 21 and day 26 iDAs following Sheta et al. Percentages show the proportion of cells positive for respective markers. **D**. iPSC-derived day 21 iDAs following Sheta et al. Cells were stained with tyrosine hydroxylase (dopaminergic neuronal marker), GIRK2 (G protein-activated inward rectifier potassium channel 2, dopaminergic neuronal marker), and DAPI (nuclear stain). Scale bar 20μm. **E**. Quantification of TH+ (over DAPI+), GIRK2+ (over DAPI+), and GIRK2+ cells that are also TH+ in day 21 iDAs following Nishimura et al. Percentages show the proportion of cells positive for respective markers. Data are presented as mean ± standard error of mean. * *p* < 0.05, ns is *not significant*

## 4. Discussion

The goal of this study was to generate a pure population of dopaminergic neurons from induced pluripotent stem cells using transcription factor-mediated differentiation. A purer population of dopaminergic neurons will allow investigations of Parkinson’s disease-associated genes and pathologies, specifically in affected dopaminergic neurons. In Parkinson’s disease, a major limitation in finding a cure is that by the time of death of the patient, 60-80% of the dopaminergic neurons are dead [25]. Therefore, it is impossible to trace back to when the dopaminergic neurons started dying. To find an early intervention to stop or halt the death of the dopaminergic neurons, we need a way of studying early mechanisms leading to death, which can’t be done in post-mortem human brain tissues. This is where a pure and functional population of iPSC-derived human dopaminergic neurons will serve as a useful model. We modified three published protocols and compared the yield of dopaminergic neurons. From our experiments, we confirm that all three protocols can generate human dopaminergic neurons; however, the yield varied considerably from what has been reported in the published protocols. We made modifications to the original protocols following Fernandopulle et al [18], for example., integration of the transcription factors in the AAVS1 safe harbour locus of iPSCs instead of using lentiviral transduction, and using human transcription factors instead of mouse transcription factors, to ensure purer population of human dopaminergic neurons. The modifications made in this study could have led to purer differentiation of iPSCs into dopaminergic neuron lineage only, but may have compromised the yield of dopaminergic neurons, particularly while following Theka et al [5] and Nishimura et al [8]. The limitation stated by Sheta et al in their protocol [26] for using lentiviral transduction was also addressed in our study by the directed and stable integration of NGN2 into the AAVS1 safe harbour locus of iPSCs. The induction step in our protocol (prior to the differentiation stage) enabled us to include a pause-step where the NGN2 induced partially differentiated cells can be cryopreserved and re-plated later. As demonstrated by Fernandopulle *et al. [18]*, the induction step provides partial differentiation, therefore, ensuring that our starting cell population for differentiation has a higher percentage of neuron-like cells and excludes any undifferentiated iPSCs. The inclusion of Ara-C from Sheta *et al*. [9, 26] also ensures that the cell population for differentiation doesn’t include any undifferentiated iPSCs. These modifications to the protocol published by Sheta et al [9] produced a better yield (> 80% TH positive and >90 % TH and GIRK2 positive) of iPSC-derived dopaminergic neurons compared to previously published protocols [5–12, 14–16] in only three weeks. Two recently published protocols [27, 28], using small molecules, succeeded in obtaining pure populations of dopaminergic neurons (> 80%) after more than a month, which were suitable for transplantation purposes. Therefore, perhaps combining the small molecule approach with transcription factor(s) will allow us to get close to 100 % yield of pure dopaminergic neurons. Finally, to test whether that the iDAs are electrophysiologically active, our GCaMP7s calcium signalling experiment showed changes in calcium flux without any stimulation.

Obtaining a pure population of dopaminergic neurons is a pre-requisite for cell replacement therapy in Parkinson’s disease. Additionally, a pure dopaminergic neuron population would help to study any Parkinson’s disease-associated gene or cellular pathology in the population of neurons that are specifically affected in the disease. However, in the human brain these dopaminergic neurons do not function alone. The neurons are supported by other brain cells such as astrocytes and microglia. As seen from our study, the percentage of TH positive neurons decreased over time and Otero et al [29] showed that healthy astrocytes helped the survival of the dopaminergic neurons. Therefore, in future experiments, combining a co-culture of astrocytes and dopaminergic neurons may facilitate better survival of the dopaminergic neurons and offer a more accurate model to study Parkinson’s disease-associated genes and pathologies. Co-culturing with other brain cells maybe be beneficial for the health of the dopaminergic neurons but will lead to an impure population of dopaminergic neurons. Therefore, once the dopaminergic neurons are mature, healthy, and functional in the co-culture system, they would need sorting (with a dopaminergic neuronal marker) to obtain a pure population. However, often sorting is harsh on the neurons and leads to cell death. Hence, maintaining a high quantity and high quality of dopaminergic neurons is one of the limitations of studies involving generation of iPSC-derived dopaminergic neurons. As transplantation requires large numbers of pure, healthy, and functional dopaminergic neurons, more optimization involving co-culture is still required to produce transplantation-grade human dopaminergic neurons.

---

**Figure S1.** Lentivirus transduction of human transcription factors in iPSCs. **A**. Two lentivirus transgene plasmid constructs were used to produce separate lentiviruses for iPSC transduction. Left map shows the plasmid carrying rtTA (reverse tetracycline-controlled transactivator) and right map shows the hALAN cassette. **B**. Both the lentiviruses were transduced in the iPSCs and FACS was used to isolate the BFP+ and neonGreen+ cells. 13.4 % cells showed double positive fluorescence. **Ci**. Post-FACS the iPSCs were positive for both markers (BFP and neonGreen), however the neonGreen was lost with passaging (**Cii**). Scale bar 100 μm.

**Figure S2.** Workflow to generate iPSC-derived iDAs following Theka *et al*. and Nishimura *et al*. **A**. The hALAN cassette was integrated into iPSCs using CRISPR-Cas9 transfection followed by expansion of the transfected iPSC. **B**. FACS was used to isolate the mApple+ iPSCs. **C**. Serial dilution, manual picking and puromycin selection were used to get pure clones of iPSCs with integrated hALAN. **D**. The selected clones were expanded and clones were validated by genotyping. **E**. Schematic of the modified iPSC-derived iDA protocol with media composition adapted from Theka *et al*. **F** and **G**. iPSC-derived day 21 and day 28 iDAs, respectively, following Theka *et al*. grown on poly-L-ornithine coated plates. Cells were stained with TUJ1 (class III β tubulin, neuronal marker), tyrosine hydroxylase (dopaminergic neuronal marker) and DAPI (nuclear stain). Scale bar 100 μm.

**Figure S3.** Differentiation of iPSCs into iDAs following Nishimura et al. **A**. Schematic of the modified iPSC-derived iDA protocol with media composition adopted from Nishimura et al. **B – E**. iPSC-derived day 21 iDAs following Nishimura et al. Cells were stained with TUJ1 (class III β tubulin, neuronal marker), tyrosine hydroxylase (dopaminergic neuronal marker), NURR1 (nuclear receptor subfamily 4 group A member 2, regulates tyrosine hydroxylase expression), LMX1A (LIM homeobox transcription factor 1 alpha, regulates the development of midbrain dopaminergic neurons), vGLUT1 (vesicular glutamate transporter 1, glutamatergic neuronal marker), MAP2 (microtubule-associated protein, 2 neuronal marker), and DAPI (nuclear stain). Scale bar 100 μm for B-D, 20 μm for E.

**Table S1.** Sequences of sgRNAs and primers used in the study.

**Table S2:** List of primary and secondary antibodies used in the study.

**Table S3:** Media composition for the modified protocol following Sheta *et al*.

## Supplementary Materials

The following supporting information can be downloaded at: www.mdpi.com/xxx/s1, Figure S1. Lentiviral transduction of human transcription factors in iPSCs, Figure S2. Workflow to generate iPSC-derived iDAs following Theka et al and Nishimura et al, Figure S3. Differentiation of iPSCs into iDAs following Nishimura et al; Table S1. Sequences, Table S2: antibodies, Table S3: Media composition; Video S1: Calcium imaging in iPSC-derived i3Ns, Video S2: Calcium imaging in iPSC-derived iDAs.

## Author Contributions

Conceptualization, I.B. and S.M.H.; methodology, K.O.M., N.L., L.S., S.G., I.B.; software, I.B., K.O.M.; validation, I.B. and L.S.; formal analysis, I.B.; investigation, I.B.; resources, I.B., S.M.H.; data curation, I.B.; writing—original draft preparation, I.B.; writing—review and editing, K.O.M., N.L., L.S., S.G., S.M.H, I.B.; visualization, I.B., L.S., N.L., K.O.M.; supervision, I.B.; project administration, I.B.; funding acquisition, I.B. All authors have read and agreed to the published version of the manuscript.

## Institutional Review Board Statement

This study was conducted according to the guidelines of the Declaration of Helsinki and approved by the University of Otago Ethics Committee (application code: APP201858, approval number: GMC100228, and date of approval: 2 March 2015) for the import and use of genetically modified iPSCs. The iPSCs were obtained from Dr. Michael Ward at the National Institute of Health, USA.

## Informed Consent Statement

No consent required as the study did not involve humans.

## Data Availability Statement

All Supplementary Materials include all the data generated in this study leading to the manuscript.

## Acknowledgments

We would like to thank the Dr. Michael Ward from the NIH for the WTC11-NGN2 iPSCs. psPAX2 was a gift from Didier Trono, Tet-O-FUW-Ascl1 was a gift from Marius Wernig, pLV.PGK.mLmx1a and Nurr1were gifts from Malin Parmar, FUW-M2rtTA was a gift from Rudolf Jaenisch, pUCM-AAVS1-TO-hNGN2 was a gift from Michael Ward, pCE-mp53DD was a gift from Shinya Yamanaka and pGP-AAV-syn-jGCaMP7s-WPRE was a gift from Douglas Kim & GENIE Project. We thank Sofia Gray for the technical support.

