## Supplementary figures and images for "Transcription factor-mediated generation of dopaminergic neurons from human iPSCs – a comparison of methods"

### Fig. S1

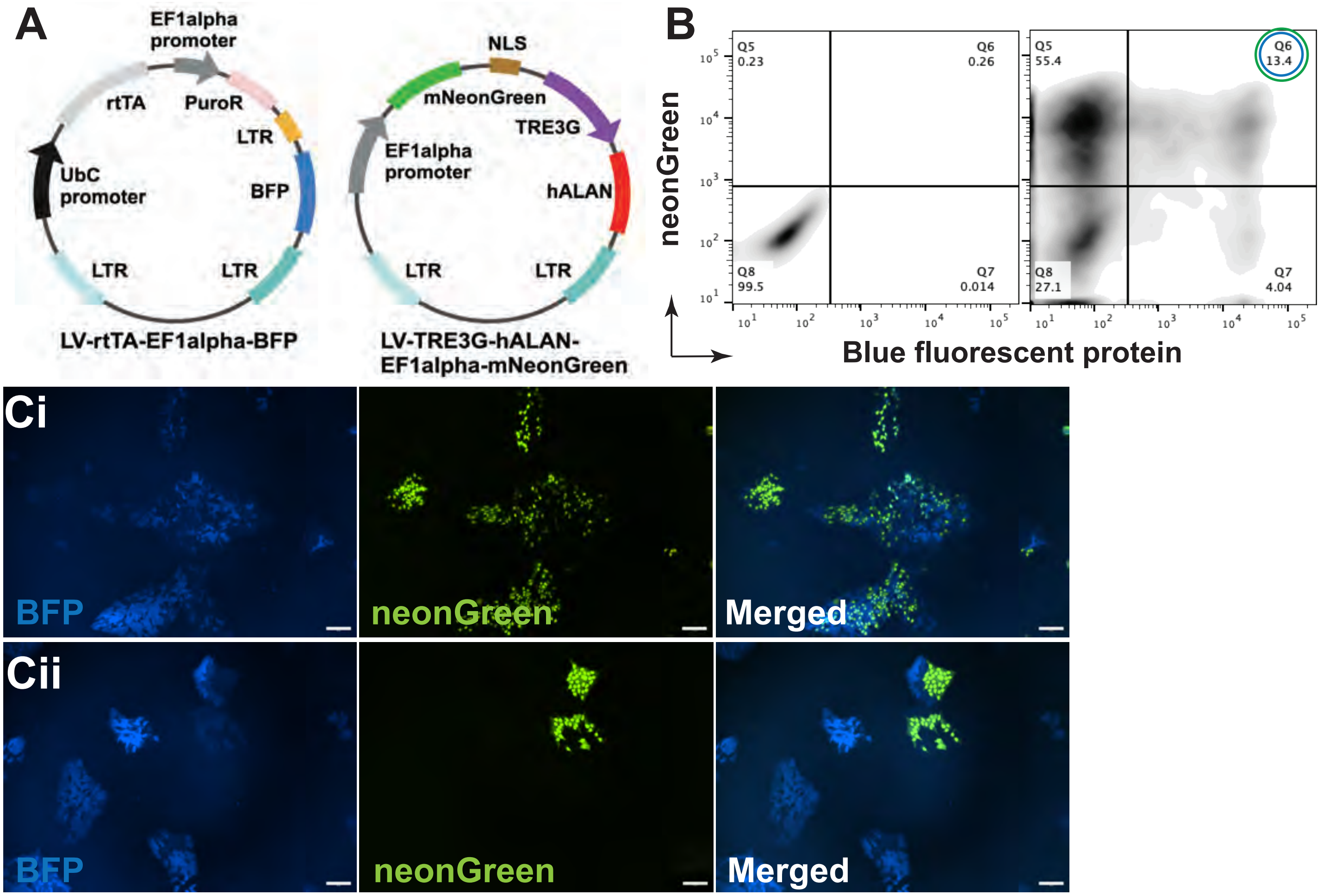

### Fig. S2

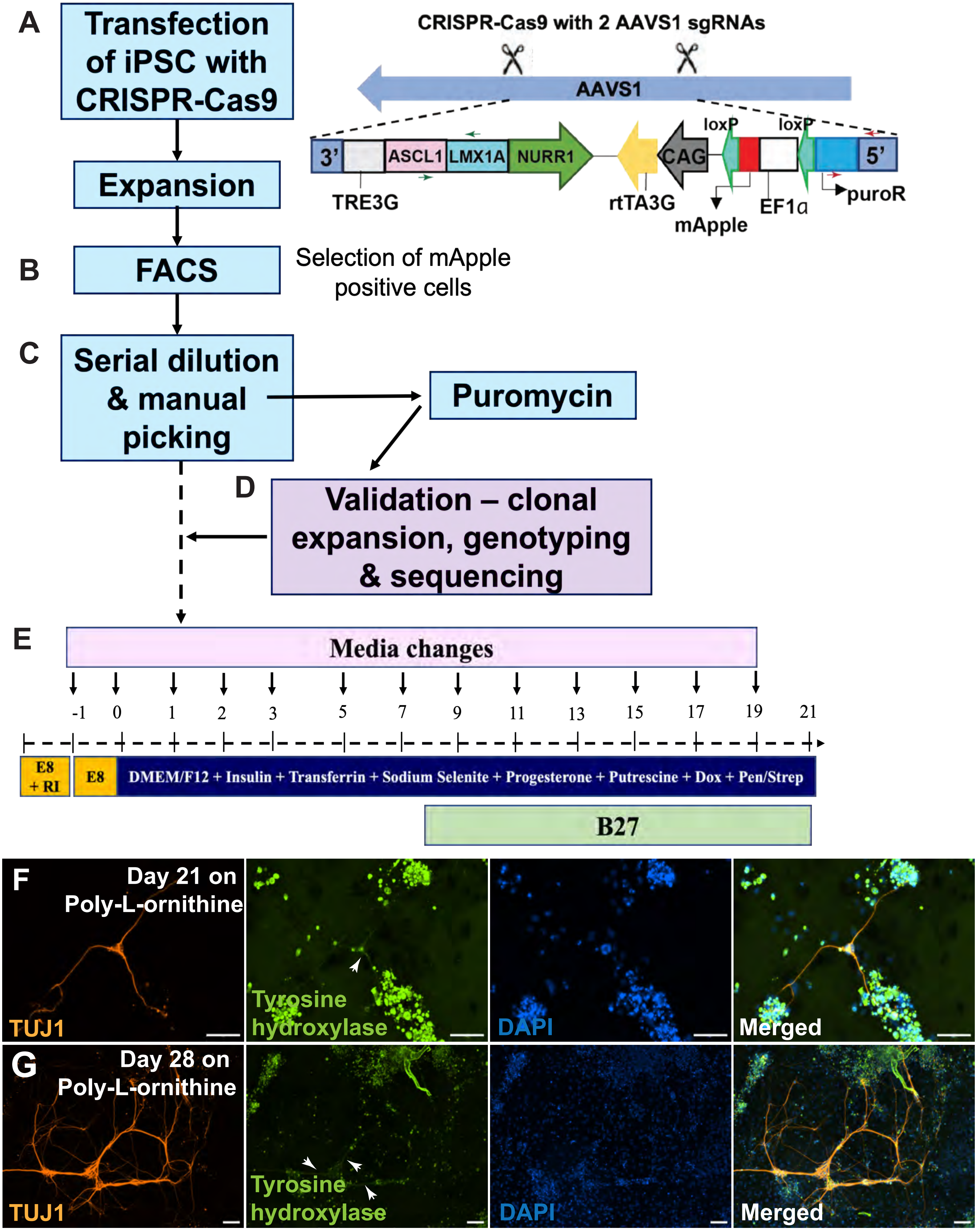

### Fig. S3

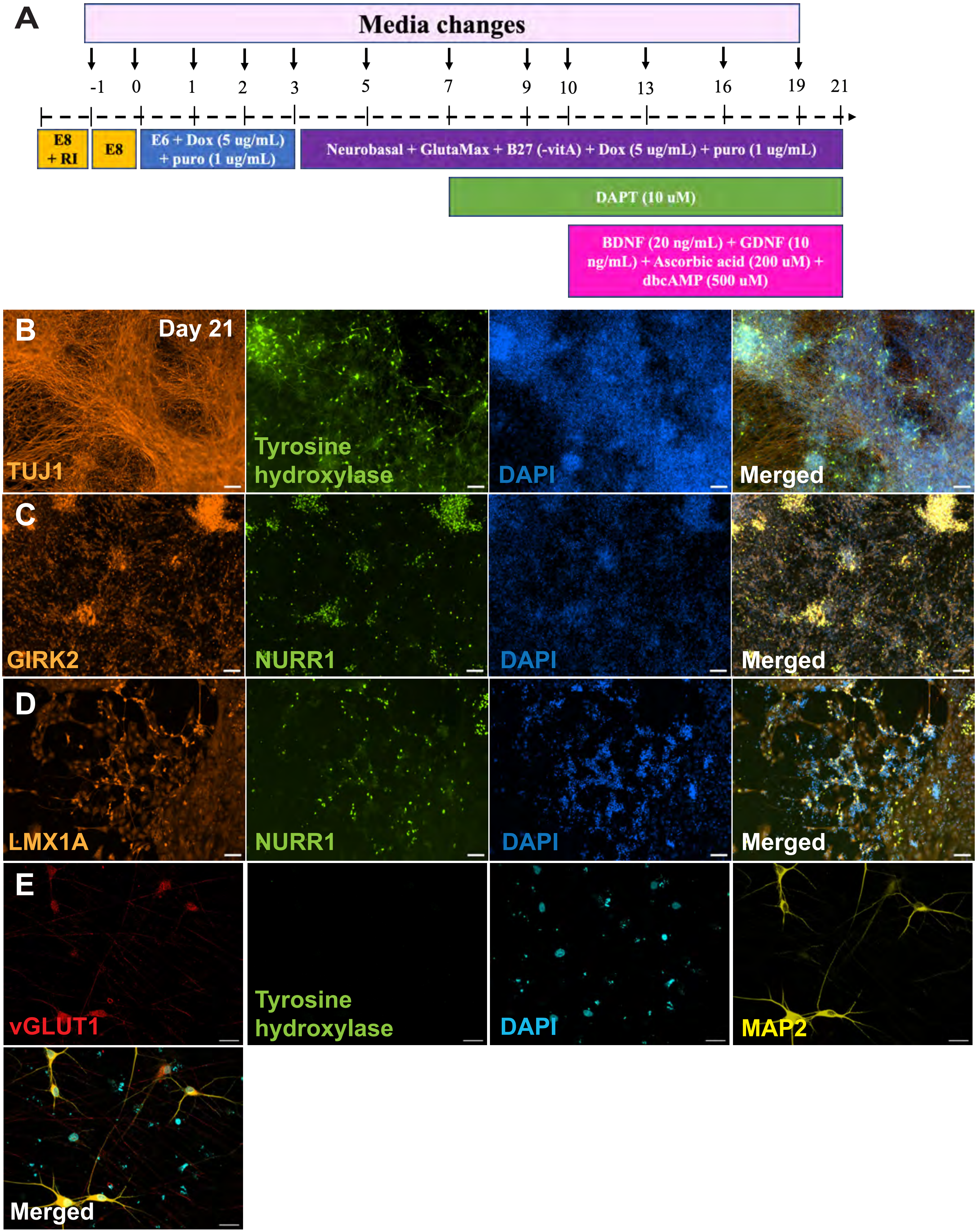
